# MaxQuant software for ion mobility enhanced shotgun proteomics

**DOI:** 10.1101/651760

**Authors:** Nikita Prianichnikov, Heiner Koch, Scarlet Koch, Markus Lubeck, Raphael Heilig, Sven Brehmer, Roman Fischer, Jürgen Cox

## Abstract

Ion mobility can add a dimension to LC-MS based shotgun proteomics which has the potential to boost proteome coverage, quantification accuracy and dynamic range. Required for this is suitable software that extracts the information contained in the four-dimensional (4D) data space spanned by m/z, retention time, ion mobility and signal intensity. Here we describe the ion mobility enhanced MaxQuant software, which utilizes the added data dimension. It offers an end to end computational workflow for the identification and quantification of peptides, proteins and posttranslational modification sites in LC-IMS-MS/MS shotgun proteomics data. We apply it to trapped ion mobility spectrometry (TIMS) coupled to a quadrupole time-of-flight (QTOF) analyzer. A highly parallelizable 4D feature detection algorithm extracts peaks which are assembled to isotope patterns. Masses are recalibrated with a non-linear m/z, retention time, ion mobility and signal intensity dependent model, based on peptides from the sample. A new matching between runs (MBR) algorithm that utilizes collisional cross section (CCS) values of MS1 features in the matching process significantly gains specificity from the extra dimension. Prerequisite for using CCS values in MBR is a relative alignment of the ion mobility values between the runs. The missing value problem in protein quantification over many samples is greatly reduced by CCS aware MBR.MS1 level label-free quantification is also implemented which proves to be highly precise and accurate on a benchmark dataset with known ground truth. MaxQuant for LC-IMS-MS/MS is part of the basic MaxQuant release and can be downloaded from http://maxquant.org.

## INTRODUCTION

Ion mobility spectrometry (1–3) (IMS) separates molecules in the gas phase by their collisional cross section (CCS) which is the effective area of a molecule quantifying the likelihood of scattering events with the gas. It can be coupled to mass spectrometry (MS) for which it constitutes a separation dimension in addition to mass over charge (m/z).Together with liquid chromatography (LC) and tandem mass spectrometry (MS/MS) one obtains LC-IMS-MS/MS shotgun proteomics, a promising strategy for the analysis of complex samples (4–7).Specifically, the object of our studies is the timsTOF Pro instrument (8) which is a time-of-flight (TOF) mass spectrometer utilizing a trapped ion mobility spectrometry (9–11) (TIMS)device operated with the parallel accumulation-serial fragmentation (PASEF) scan mode (12).

MaxQuant (13, 14) is a popular software platform for LC-MS/MS shotgun proteomics possessing a large ecosystem of algorithms for comprehensive data analysis (15). It incorporates the peptide search engine Andromeda (16) and the companion software Perseus (17, 18) offers a complete solution for the downstream bioinformatics analysis. MaxQuant performs quantification with labels (19) and via the MaxLFQ algorithm (20)on label-free data. MaxQuant achieves high peptide mass accuracies thanks to its advanced nonlinear recalibration algorithms (21, 22). It contains comfortable visualization capabilities (23) for the inspection of the raw data and runs on Windows and Linux operating systems (24).

The aim of this publication is to combine these two factors, i.e. to make MaxQuant capable of analyzing timsTOF Pro data. A main challenge originates from the inflation of raw data by the added dimension. To keep the computation time required for the analysis down to a realistic amount is of highest importance. In this manuscript we describe the updated computational workflow of MaxQuant for ion mobility-enhanced shotgun proteomics data, which has been optimized for computational performance. It is based on 4D feature detection in the space augmented by the extra dimension provided by IMS. Matching between runs is a crucial algorithmic step in MaxQuant for retrieving MS1 features in data-dependent acquisition shotgun proteomics, in order to make quantification of peptides and proteins more reproducible over many samples. It requires high mass accuracy and precise relative retention time measures, obtained by non-linear retention time alignment in order to restrict false positives matches. Here we present a new matching algorithm that takes into account ion mobility values of peptide features measured on a timsTOF Pro instrument. Similarly, as for the retention times, we observe the necessity for relative alignment of ion mobility values between LC-IMS-MS/MS runs, and include a nonlinear multi-sample alignment algorithm into the MaxQuant workflow. Matching the 4D features using these aligned CCS values did significantly improve specificity. Furthermore, we observe a strong positive impact on the missing value problem in quantitative proteomics. All new algorithmic steps as well as their application to example data are detailed in the Results section.

## EXPERIMENTAL PROCEDURES

### Cell culture and sample preparation

Whole protein extracts of human cervical cancer cells (HeLa) were purchased fromPromega and digestion was performed according to the protocol of Wang et al.(25). Briefly, the protein lyophilisate was re-constituted in water and trifluoroethanol (1:1). Disulfide bonds were reduced with dithiothreitol at a concentration of 5 mM and alkylated with chloroacetamide (20mM) in ammonium bicarbonate buffer, followed by 90 min incubation in the dark. Trypsin (Promega) was added in a protease: protein ratio (wt:wt) of 1:100 and incubation was performed overnight at 37°C. Digestion was stopped by acidification with formic acid to pH 2 and peptides were desalted on C18 cartridges (3 M Empore) and dried in a vacuum centrifuge. For the mixed species experiments, tryptic protein digests of H. sapiens (HeLa), S. cerevisiae (Promega) and E.coli (Waters) were mixed in two different experiments leading to a ratio of 1:1 (HeLa), 1:2 (S. cerevisiae) and 1:4 (E.coli) between the two samples. The human blood plasma samples were collected from acute inflammation patients and were depleted for the 12 most abundant proteins using spin columns (Pierce) according to the manufacturer’s instructions. The depleted plasma proteome was TCA/DOC precipitated and digested with SMART digest trypsin (Pierce) for 2h. Peptides were desalted (SOLAμ HRP, Pierce), diluted to 5 ng/μL0.1% of formic acid in water for subsequent transfer of 100 ng to Evotips.

### Liquid chromatography

A nanoElute (Bruker) high pressure nanoflow system was connected to the timsTOF Pro, an ion-mobility spectrometry quadrupole time of flight mass spectrometer (Bruker). Peptides were reconstituted in 0.1% FA and 200 ng were delivered to reversed phase analytical columns (25 cm x 75 µm i.d., Ion opticks) with pulled emitter tips. Liquid chromatography was performed at 50 °C and peptides were separated on the analytical column using a 120 min gradient (solvent A: 0.1% FA; solvent B: 0.1% FA, in ACN) at a flow rate of 400 nl/min. A linear gradient from 2-17 % B was applied for 60 min, followed by 17-25 % B for the next 30 min and followed by a step to 37% B for 10 min and a step to 80% B for 10 min followed by 10 min of washing at 80% B. For the plasma proteome samples, the Evosep One (Evosep) was used to achieve low overhead times and a high sample throughput. Separation was performed by transferring 100 ng proteolytic digest from the sample loop on short columns (8 cm x 100 µm i.d.) at flow rate of 1.5 µl/min achieving 11.5 min gradient times and the measurement of 100 samples/day.

### Ion mobility mass spectrometry

For all experiments, the timsTOF Pro was operated in PASEF mode. Ions entering the instrument were orthogonally deflected into an electrodynamic funnel and were trapped in the front region of the trapped ion mobility (TIMS) analyzer. The TIMS tunnel is separated into two parts (“dual TIMS” design), allowing for accumulation of entering ions in the first part and trapped ion mobility elution in the second part. The dual TIMS design allows usage of almost 100% of the ions as long as equal accumulation and analysis times are used. While ions are eluted from the second part, accumulation of new ions can already take place in the first part for subsequent transfer into the second part. Trapped ion mobility separation was achieved by repulsion of an increasing longitudinal electric field gradient (ramp) and a drag force of incoming gas from ambient air. Dependent on the collisional cross sections and charge states high mobility ions are accumulated at the front part and low mobility ions at the end of the analyzer. To achieve close to 100 % duty cycle, we set the ramp and accumulation time for the electric field gradient to 100 ms. For each topN acquisition cycle and long nanoLC runs, one full frame and 10 PASEF MS/MS frames, each containing on average 12 MS/MS spectra, were acquired resulting in a cycle time of 1.1 s. MS and MS/MS spectra were recorded from 100 to 1,700 m/z and precursor ions for PASEF scans were selected in real time by the precursor selection algorithm. A polygon filtering was applied in the m/z and ion mobility area to exclude the low m/z of singly charged ions for PASEF precursor selection. For the 2h runs a ‘target value’ of 20.000 was applied to repeatedly schedule MS precursors for PASEF MS/MS spectra until this intensity value is reached and an ion mobility range (1/*K_0_*) of 0.6-1.6 Vs/cm^2^ was used. By using one full frame and 4 PASEF MS/MS frames we have optimized data acquisition for short gradients on the Evosep system to achieve a good MS1 sampling rate for quantification at a cycle times of 0.5 s. A ‘target value’ of 6.000 was applied to repeatedly schedule MS precursors with low intensity in digests of plasma proteomes and the ion mobility range was limited to 0.85-1.3 Vs/cm^2^. For all experiments the quadrupole isolation width was set to 2 Th for m/z < 700 and 3 Th for m/z > 700. Collision energy was changed in 5 steps within a TIMS elution ramp from 52 eV for 0-19% of the ramp time; 47 eV from 19-38%; 42 eV from 38-57%; 37 eV from 57-76%; and 32 eV for 76-95%.

### MaxQuant software framework

MaxQuant version 1.6.6.0 was used to perform all data analysis, which is downloadable from http://maxquant.org. MaxQuant has a plugin API for raw data access, which has been implemented for the access to Bruker TDF raw data format. MaxQuant is written in C# (.NET Framework 4.7.2 or higher) and runs on Windows and Linux operating systems(24). MaxQuant can be run interactively from a user interface or alternatively be called from the command line. Projects analyzed in MaxQuant can be automatically uploaded to the PRIDE repository as a ‘complete’ submission(26).A MaxQuant help forum can be visited at https://groups.google.com/group/maxquant-list/. Bug reports should be addressed to https://maxquant.myjetbrains.com/youtrack/.

For searches with HeLa data we used human UniProt sequences (https://www.uniprot.org/proteomes/UP000005640, version January 26, 2019) containing 73,920 proteins. All searches were performed with oxidation of methionine and protein N-terminal acetylation. Values of parameters in MaxQuant have not been changed from their default values unless explicitly stated.

## RESULTS

### Computational workflow for data-dependent LC-IMS-MS/MS data

The MaxQuant workflow is an end-to-end solution, taking the mass spectrometric raw data as input and providing output tables on several levels, e.g., a table for results on the level of protein groups, of peptides or of peptide spectrum matches (PSMs). The identification strategy is peptide database search engine-based, utilizing the integrated Andromeda search engine as the main source of identifications. Also, for ion mobility-enhanced data the identification of peptides will be done by conventional Andromeda searches for which the input spectra are projected to the usual form of m/z-intensity profiles. The workflow consists of a series of data processing steps, the most important ones of which are summarized in **Table 1**. Some computational steps needed substantial adaptation in order to support IMS, while other steps remained nearly unchanged. For instance, feature detection needed to be massively adapted, since adding a dimension to the raw data space requires new concepts for the detection of features. Also the preparation of MS/MS spectra for database search is vastly different for ion-mobility enhanced data, since it involves a projection from the higher-dimensional data to conventional spectra. The alignment of raw data between samples will be affected since, in addition to the alignment of retention times, it may be necessary to also calibrate ion mobility measurements relative to each other. The process of matching features between runs for the sake of increasing the feature set usable for quantification will also need adaptations since the collision cross section values can be used to make the matching more specific. On the other hand there are algorithms that need no adaptation from the standard workflow: for instance, grouping proteins into redundant protein groups or determining their false discovery rate (FDR) will remain unchanged. In the following sub-sections we describe how the individual data processing steps have been adapted for LC-IMS-MS/MS data.

**Table 1.** Main processing steps of the computational workflow. The table lists the most important and time consuming steps in the MaxQuant pipeline for LC-IMS-MS/MS data. In the second column, the number of plus signs signifies the extent of changes in the code in order to accommodate ion mobility information. For instance, ‘+++’ for ‘feature detection’ indicates fundamental changes. The right column contains the percentage of computation time that is spent on the step on the HeLa dataset consisting of ten replicates, on 60 core machine running Windows Server 2016.

| Workflow component | Changes for IMS | Percentage computation time |
| --- | --- | --- |
| Feature detection | +++ | 31.2% |
| MS/MS preparation | +++ | 15.6% |
| Initial search | - | 14.1% |
| Recalibration | ++ | 1.8% |
| Main search | - | 19.1% |
| PSM FDR | - | 1.9% |
| Alignment | ++ | 12.5% |
| Match between runs | ++ | 0.4% |
| Protein groups / FDR | - | 2.6% |
| Label-free quantification | - | 0.3% |
| Writing of results | + | 0.4% |

### Feature detection

The task of feature detection becomes more challenging in the higher-dimensional ion mobility-enhanced data. While in LC-MS data, peak boundaries were determined by a closed line in the *m/z*-retention time plane, now a peak is bounded by a two-dimensional closed surface in three-dimensional space. In principle the task is similar to segmentation of 3-dimensional voxel data into regions of independent signals. However, since the shapes of feature boundaries in MS data typically exhibit certain regularities, we want to exploit these to obtain a well performing algorithm, since otherwise computation time remains a bottleneck for ion-mobility enhanced MS data. **Fig. 1** shows an overview of the steps involved in feature detection. First, the data is interpolated onto a common mass grid (**Fig. 1a**) while the number of scans in the ion mobility direction remains the same. The lattice spacing of the new *m/z* grid depends on *m/z*,to ensure that the local point density is adapted to the peak width that is approximately expected by the resolution, and is calculated such that

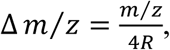

where *R* plays the role of an effective instrument resolution which we take as 32,000 by default. For interpolating intensities into one of the resulting spectra we take into account a window of ion mobility indices around the current index over which the signals are averaged. All raw intensity measurements within the ion mobility window and within a mass window corresponding to three sigma of the peak resolution *R* are smoothed with a Gaussian kernel, whose width is locally adapted to correspond to the resolution *R*. This results in a data cube with a regular gridding in all three dimensions and smoothly varying signal intensities over the cube. All the data gridding and smoothing is done on the fly when a spectrum from the data cube is needed and, in order to save space, not written to disk. We proceed by slicing the cube into planes perpendicular to the ion mobility dimension (**Fig. 1b**). For each fixed ion mobility value we obtain a pseudo LC-MS run with signal intensities depending only on m/z and retention time. In these we apply the conventional MaxQuant algorithm for detecting peak boundaries as closed lines. Neighboring ion mobility slices are usually similar to each other regarding their signal intensity distribution. Therefore, in order to save computing time, ion mobility slices can be omitted and only every *n*^th^ slice is used in the data analysis, where *n* is a user-definable parameter with default value set to three. The original MaxQuant feature detection algorithm processes then each of these slices resulting in irregularly shaped peak boundaries, which, in particular, do not have to be rectangular (**Fig. 1c**).Since the processing of one slice does not depend on the processing of other slices, this step is trivially parallelized onto a specified number of threads less or equal to the number of slices.Once the peak boundaries in the slices are obtained, overlapping areas are clustered across slices to obtain closed surfaces enclosing *m/z*-retention time-ion mobility volumes (**Fig. 1d**). These volume elements define the base of each feature and each point inside the volume has an intensity value attached, defining the m/z, ion mobility and retention time-dependent intensity profile of the feature. The total feature intensity is given by the integral of signals over the volume.

**Figure 1.**
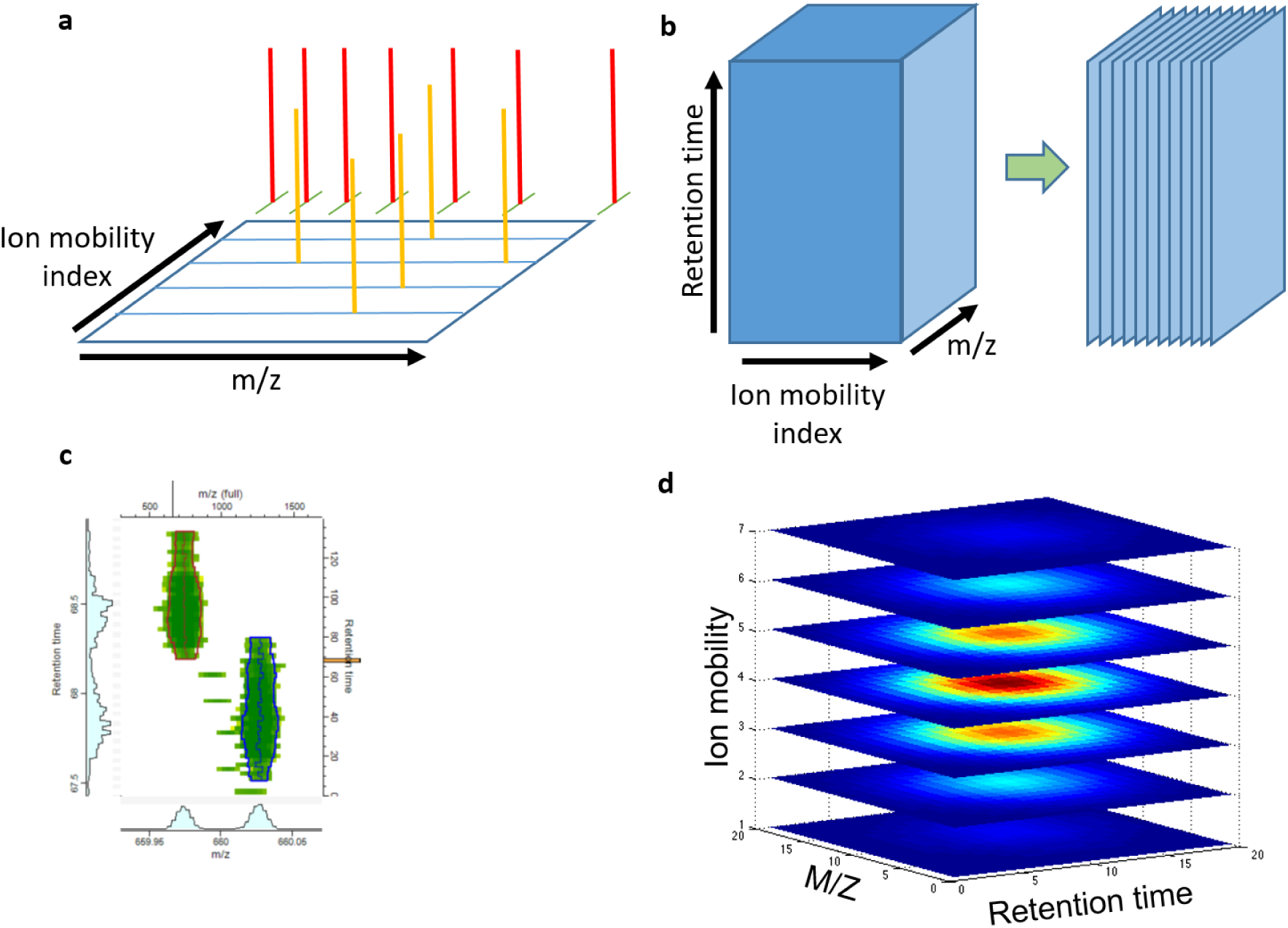
Elements of feature detection. **a.** Raw data is mapped to a common grid. While the ion mobility scans already form a regular grid, the *m*/*z* values of data centroids are irregular. All intensity values within an ion mobility window and within a mass range are mapped to a common mass grid with a spacing that is monotone increasing with m/z. Intensities are added up using a Gaussian kernel with a locally adapted with according to the effective resolution. **b.** The raw data is sliced along the ion mobility axis to obtain data planes with signal intensity as a function of *m*/*z* and retention time. In these planes, features can be detected with the conventional algorithms in MaxQuant. **c.** The features in each plane are bounded by irregular shapes following the raw data. **d.** Features are clustered between consecutive planes to obtain closed surfaces surrounding the final features.

The de-isotoping step aims at grouping those peaks, which are different isotopic forms of the same peptide molecule, together into isotope patterns. For this purpose we generalized the de-isotoping algorithm in MaxQuant, which makes use of the correlation of peak intensities as a function of retention time, to also take into account correlations in ion mobility direction. The isotopic peak clustering is a two-step process. The first step consists of a pre-clustering of features. Here, two features are put together into a cluster whenever two criteria are met: 1) their difference in m/z is compatible with an averagine (27) mass difference between two consecutive peaks in an isotope pattern and 2) The cosine correlation between the two intensity patterns over retention time and ion mobility exceeds a threshold. Since these pre-clusters are assembled based on pair-wise relations between peaks, they can be too large and inconsistent regarding charge state, typically containing, in addition to the main isotope pattern of the cluster, other correlating peaks. These are separated in the second step which refines the pre-clusters based on several criteria, as, for instance, consistency of charge state. As an example we report numbers of scans, features, isotope patterns and identifications found in a single typical LC-IMS-MS/MS run in **Table 2**.

**Table 2.** Numbers of features and other elements in a single LC-IMS-MS/MS run. For a typical run with a two-hour gradient we show the numbers of scans, detected elements and identified entities.

| **Workflow component** | **Number of items per run in HeLa sample** | **Number of items per run in plasma sample** |
| --- | --- | --- |
| **MS1 frames** | 7828 | 1729 |
| **MS2 frames** | 52679 | 3990 |
| **IMS scans per frame** | 918 | 918 |
| **PASEF precursors** | 233437 | 44214 |
| **4D features** | 2164978 | 540917 |
| **4D isotope patterns** | 409379 | 65501 |
| **MS/MS submitted** | 114477 | 16928 |

### Extraction of MS/MS spectra

The timsTOF Pro effectively gains MS/MS spectrum acquisition speed by measuring multiple PASEF fragmentation patterns in one frame. It generates up to 110 of these MS/MS frames per second, which, in the example shown in **Table 2** amounts to 233437 raw PASEF precursors distributed over 52679 MS/MS frames in a two-hour LC-IMS-MS/MS run. Multiple scans can be triggered for the same precursor. This is done deliberately, because the acquisition software decides to allocate additional MS/MS scans to a low-abundant precursor in order to accumulate signal for the fragmentation spectrum, to increase the likelihood of its identification. There can be additional multiplicity in terms of multiple MS/MS scans per precursor, which happens accidentally, and only becomes visible post acquisition by the MaxQuant analysis. These resequencing events are usually further apart in retention time and only the 4D peak detection reveals that they belong to the same precursor. All MS/MS spectra that were acquired for the same precursor are accumulated in MaxQuant. In them equal ions are added up, and only a single summed spectrum per 4D precursor is submitted to the database search. Fragmentation spectra are projected from the higher-dimensional raw data by integrating the MS/MS frame intensities over the ion mobility index range for each PASEF scan. This results in a single conventional MS/MS spectrum per MaxQuant precursor that can be submitted to the Andromeda search engine in the conventional way.In the projected and summed spectra de-isotoping is performed and in case an isotope pattern is detected, its members are removed from the spectrum and added as the monoisotopic peak of a singly charged fragment. If this procedure leads to overlapping peaks within a mass tolerance, resulting from multiple charge states for the same fragment, the corresponding peak intensities are added up to a single peak.

In the example run in **Table 2** the initial number of PASEF precursors is 233437 (run 20190122_HeLa_QC_Slot1-47_1_3219.d) which gets accumulated to 114477 MS/MS scans by MaxQuant corresponding to unique 4D precursors. Only these accumulated scans are submitted to the database search. Usually, a large fraction of MS1 precursors remains without an MS/MS scan taken for them (294,902 in the example in **Table 2**), which makes them ideal targets for exploiting them for quantification with the help of the ion mobility-enhanced matching between runs algorithm. (See later subsection on alignment and matching between runs.)

### Mass recalibration

Nonlinear mass recalibration is a crucial prerequisite for obtaining high peptide mass accuracies in MS-based proteomics. For LC-MS/MS data, MaxQuant performs nonlinear recalibration of the m/z range. The recalibration function used for this depends nonlinearly on m/z, retention time and signal intensity. Peptides present in the sample are used as standards, which eliminates the requirement for spike-in of molecules as calibration standards. A preliminary round of peptide identification with simple identification criteria, as, for instance, a fixed Andromeda score cutoff, is performed for generating such a list of peptides that are going to be used as internal standards. By default we use an Andromeda score cutoff of 70. Once the recalibration function is determined and applied to the whole data, in the subsequent analysis, more stringent mass tolerances can be applied to precursor masses, for instance in the main peptide search and in the matching of MS1 features between runs. These precursor mass windows can even be adapted to each individual peptide by exploiting the variability of multiple mass measurements within the 4d precursor. Also, in the main search more refined statistical methods will be used for the peptide identification process than a simple score cutoff.

Ion mobility adds another variable, potentially confounding the mass error of MS1 features in a nonlinear way. Hence, the m/z recalibration function can now depend on four variables in total. We split the general dependence of the mass error in parts per million (p.p.m.) into four additive components, each of them depending only on a single variable

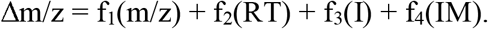

A constant contribution can be arbitrarily split among the four contributions. Hence, three of the four functions can be constrained to have a certain value, e.g. at an argument of choice. We determine first the ion mobility dependent part f_4_(IM) by calculating a window median of the mass deviations over ion mobility index windows and subtract it from the original mass deviations. We then apply the conventional MaxQuant algorithm depending on the other three dimensions to the ion mobility independent residual deviation. The m/z and retention time dependent parts are modeled as piecewise linear functions, while the intensity dependent part is assumed to be a low degree polynomial in the logarithm of the signal intensity.

In **Fig. 2a-d** we show the individual contributions of the four variables to the mass error. For each variable, the residual error is shown after the other three dependencies have been taken into account. For instance in **Fig. 2a** Δm/z - f_2_(RT) - f_3_(I) - f_4_(IM) is plotted against m/z to show the m/z dependent contribution to the mass error. It can be observed that the ion mobility-dependent component has the largest amplitude. It explains 52%of the variance captured by the model. Hence, it is crucial to include an ion mobility-dependent component in the mass recalibration. The residuals after complete recalibration (Supplementary Fig. 1) show no remaining systematic effects. As can be seen in the mass error histograms before and after recalibration (**Fig. 2e-f**) the mass accuracy improves considerably upon recalibration. The median absolute deviation (MAD) is 2.79 p.p.m. before and 0.94 p.p.m after recalibration, corresponding to a3-fold improvement.

**Figure 2.**
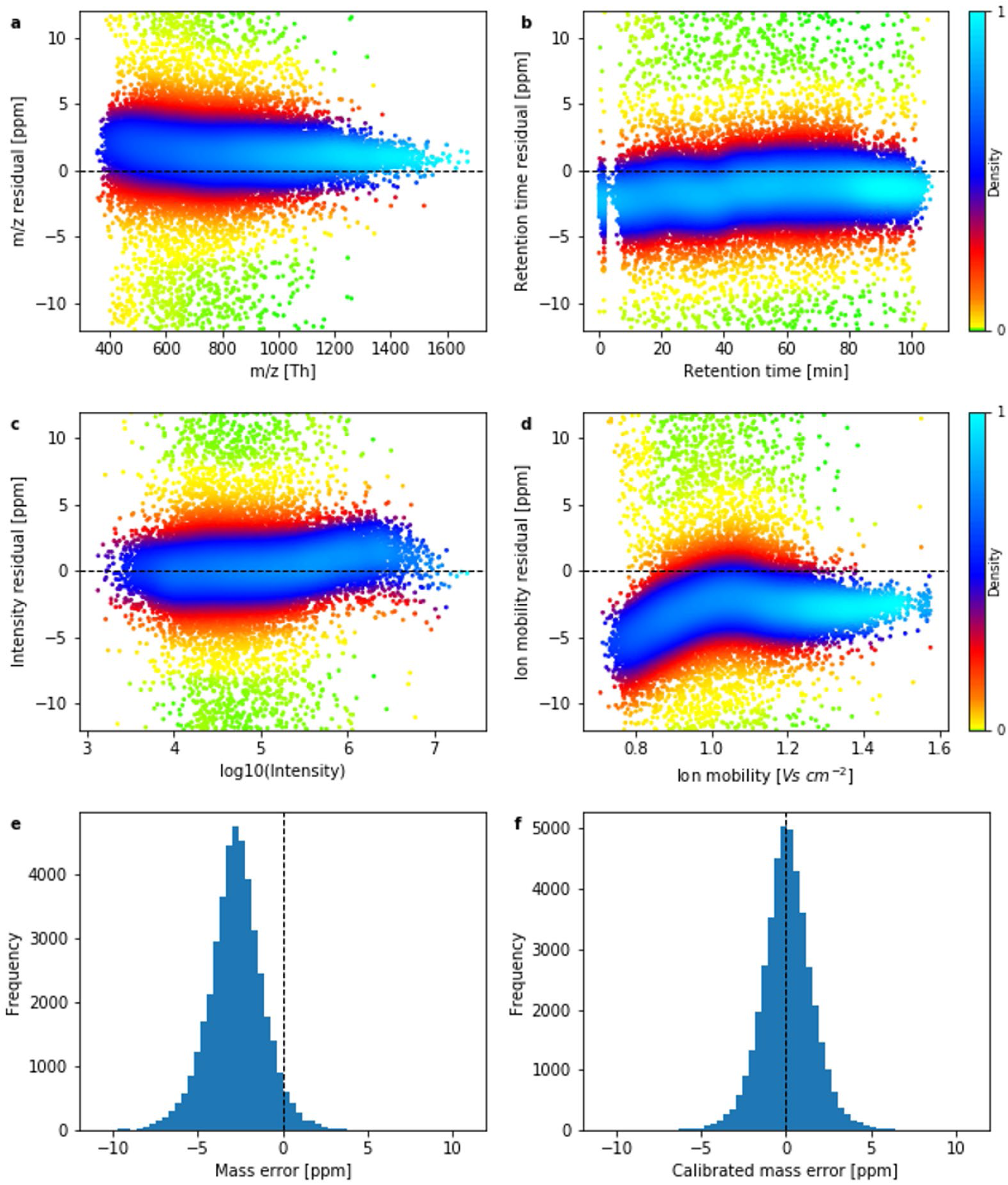
Mass recalibration. **a.-d.** Residual mass errors after the dependence of all but one variable have been recalibrated, showing the dependence of the residual mass error on *m*/*z* (**a.**), retention time (**b.**), logarithm of the peak intensity (**c.**) and ion mobility (**d.**). Colors reflect the density of data points. **e.** Mass error distribution before recalibration. **f.** Mass error distribution after recalibration has been applied.

### Alignment and matching between runs

To alleviate the stochasticity of shotgun proteomics, MaxQuant employs the matching between runs algorithm, which consists of transferring identifications of MS1 features between samples based on accurate mass and retention time values (28). One promise of LC-IMS-MS is that this matching between runs becomes more specific by exploiting the additional dimension of ion mobility, which can be used to eliminate false positive matches. In order to use the retention time as a criterion for matching features, MaxQuant allows to first perform a nonlinear recalibration of retention times. It is an open question if such an alignment is also necessary prior to using ion mobility values for this kind of matching.

To address this question we matched features between two LC-IMS-MS/MS runs. These were runs 2823 and 2799 from the dataset ‘20181129HeLafraction22’, which were measured on the same mass spectrometer and column, but days apart without recalibrating mass or mobility. We first find feature pairs with one feature from each run by applying mass, retention time and ion mobility windows. Masses are already recalibrated, which implies that small windows can be used. The actual criterion for matching features between the two runs applies the individual mass tolerances determined by MaxQuant after mass recalibration. Furthermore we use wide windows for retention time and ion mobility index since the alignment has not yet been performed in these two dimensions. For instance, time windows of several (15 by default) minutes could be used. Whenever there is more than one matching feature found in the second run for a feature in the first run, we take only the closest match according to a weighted distance in m/z, retention time and ion mobility index space.

Based on the feature pairs found by this procedure, we create plots of retention time difference within each feature pair against retention time in one of the runs (**Fig. 3a**) and similarly for ion mobility index difference against ion mobility index (**Fig. 3c**). In each plot the local point density has been color coded. As can be seen for this typical case, a nonlinear recalibration function needs to be applied in order to make the retention times in the two runs comparable. This is also the case for the ion mobility index: it is necessary to do ion mobility alignment in addition to retention time alignment. In the example shown, a linear recalibration would be sufficient. Nevertheless, we apply the same nonlinear recalibration algorithm that is used for retention time alignment also for the ion mobility direction, in order to be prepared for the general case. As can be seen in **Fig. 3b** and **d**, after alignment the retention times and ion mobility indices become comparable between runs. After alignment, the matching requirement is, in addition to the masses matching, that the peak maximum is within a tighter retention time window, typically 42 seconds, and the aligned ion mobility index match is bounded by a tighter window as well.

**Figure 3.**
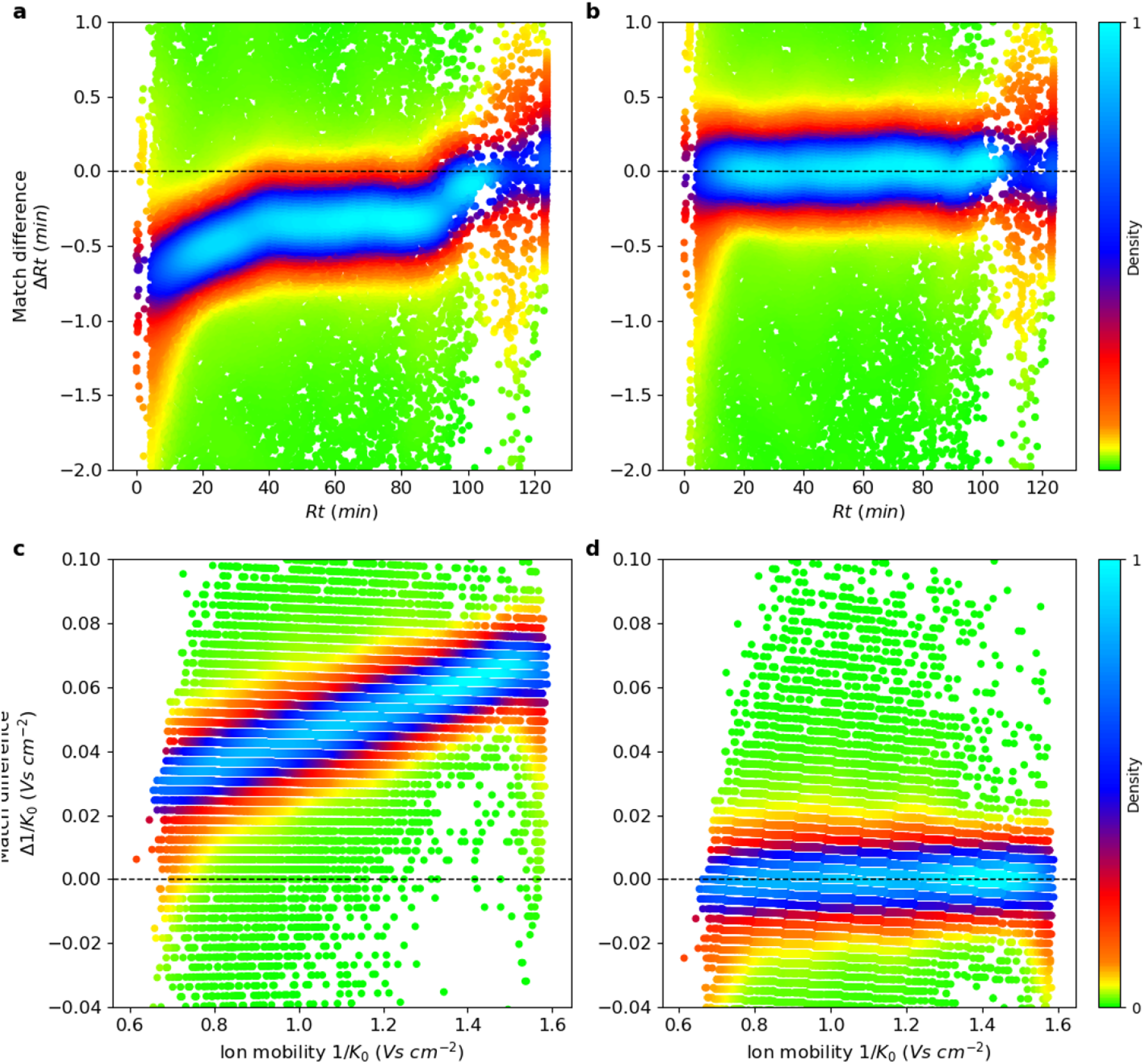
Retention time and ion mobility alignment. **a.** Difference in retention time between matched feature pairs plotted against the retention time of the feature in one of the runs. The point density is color-coded in plots **a-d**. **b.** Same as in **a**, but after retention time alignment has been applied. **c.** Similar to **a**, but now the difference in 1/*K_0_* within the feature pair is plotted against 1/*K_0_* in one of the runs, indicating differences between runs in terms of ion mobility. **d.** Same as in **c**, but after ion mobility alignment has been applied.

The distribution of time differences between matching features as well as of the ion mobility index differences are shown in **Fig. 4**.In **Fig. 4a** the matching time differences before alignment are shown while **Fig. 4b** displays the distribution of time differences after alignment. Only after alignment is the distribution centered on zero. The full width half maximum (FWHM) for the retention time differences is 8.5 seconds after alignment in this two-hour run. **Fig. 4c-d** shows similar distributions before and after alignment for the ion mobility index. As can be seen, besides a centering to zero, the FWHM improves dramatically through alignment, decreasing by a factor of 2.59, which brings it down to less than 1% of the total ion mobility range. This again displays the need for relative alignment of ion mobility measurements in the general case before it is being used for matching of features. Aligned and calibrated ion mobilities should then be comparable between different platforms, since they are a property of the molecule and not of the measuring systems, as retention times are.

**Figure 4.**
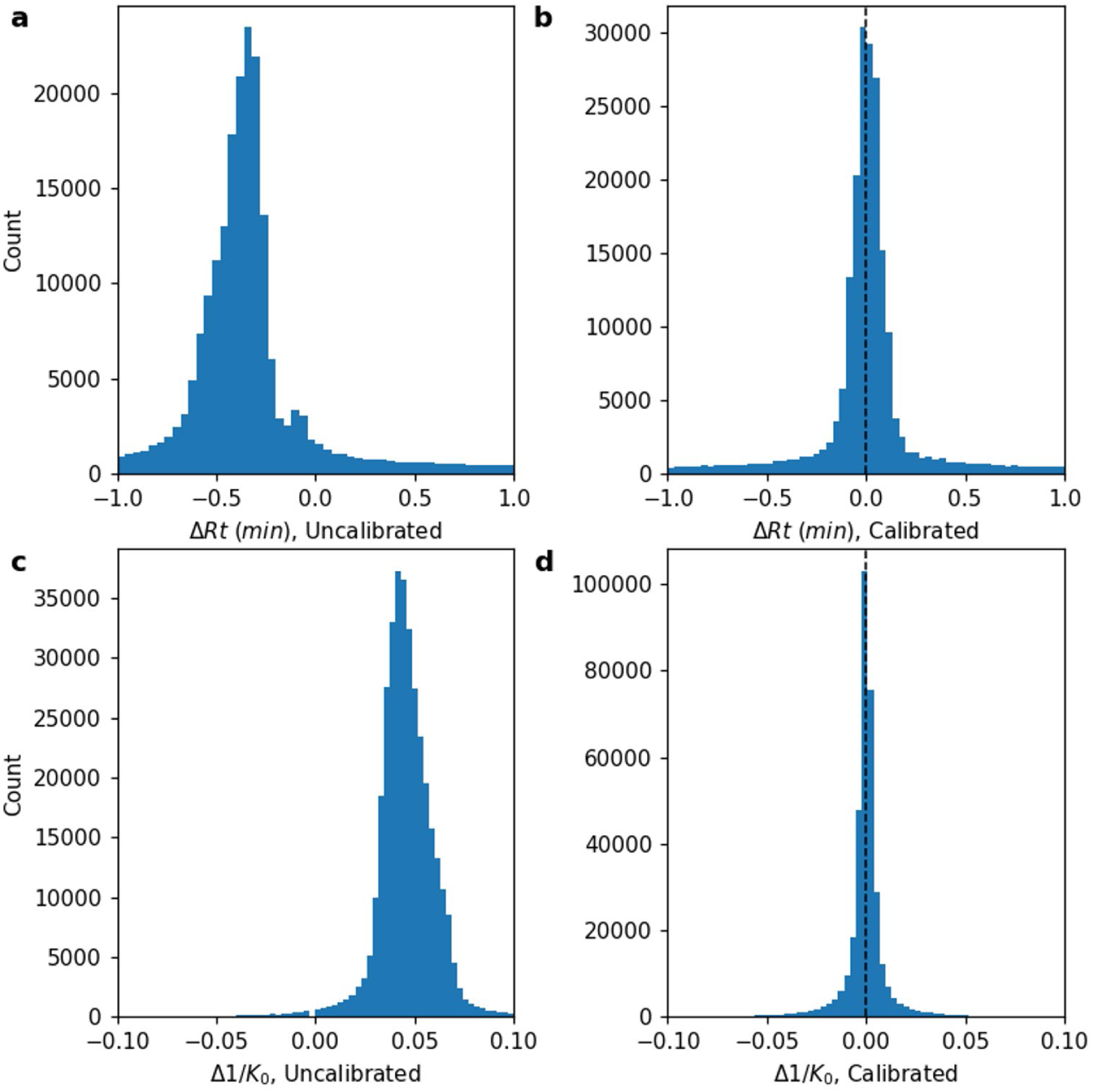
Accuracy of matching between runs. **a.** Retention time match difference distribution before alignment. **b.** Retention time match difference distribution after alignment. **c.** 1/*K_0_* match difference distribution before alignment. **b.** 1/*K_0_* match difference distribution after alignment.

While so far we have described how two runs are aligned and its features are matched, MaxQuant can perform the same for multiple samples as well, without singling out one of the samples as a master run. The alignment is done by first building a guide tree based on the similarity between the runs. The alignment starts then between the two most similar runs. After these have been aligned they are merged to an effective aligned sample. The process continues by always taking the as yet not merged most similar samples from the guide tree, align and merge them, and continue until all samples are aligned.

In order to estimate the gain in accuracy that is brought about by using ion mobility as an additional criterion for matching features between runs, we perform a special analysis of matches with relatively loose criteria. Now, matches are performed without using retention time but with applying a varying matching window in 1/*K_0_*. (See **Fig. 5**.)Then we want to use the fact that matches with large retention time differences have a larger likelihood of being false than matches with small retention time difference. We introduce the (not strictly correct) nomenclature of calling a match with retention time difference larger than 42 seconds a ‘false’ match and when it is smaller than 42 seconds a ‘true’ match. Note that the threshold is picked arbitrarily, just to allow us distinguishing two populations of matches one of which is enriched and one depleted of false positives. In **Fig. 5a** the number of ‘true’ and ‘false’ matches is shown as a function of the window size in 1/*K_0_* used for accepting matches. Both curves are plateauing to the right which corresponds to only mass-based matching. When decreasing the 1/*K_0_* window the ‘false’ matches are decreasing more rapidly than the ‘true’ matches. In order to quantify the 1/K0 window dependent gain we show in **Fig. 5B** the ratio of the two curves for ‘true’ and ‘false matches, which before are normalized to both plateau at 1. While a large 1/K0 window provides no gain in terms of specificity, a smaller window provides an up to three-fold gain in specificity.

**Figure 5.**
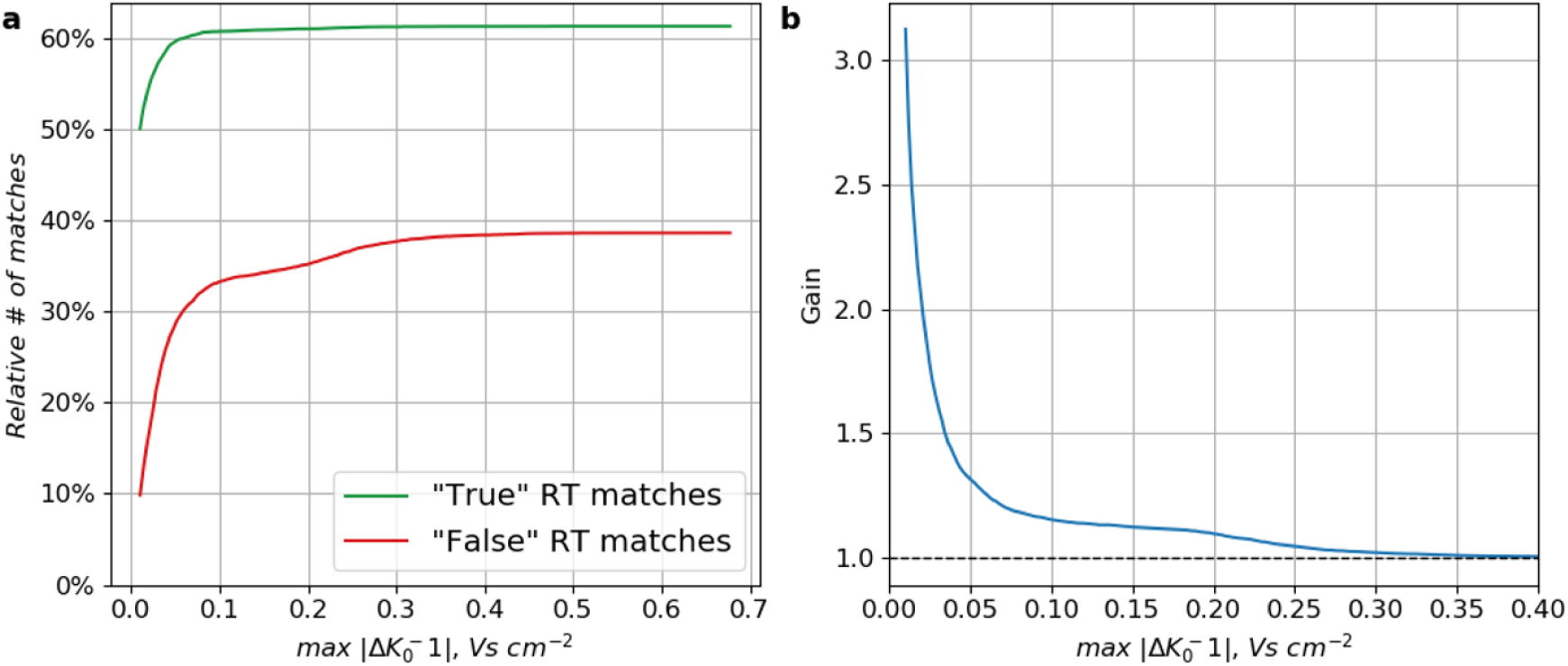
Specificity of ion mobility enhanced matching between runs. **a.** Matches were performed without retention time restriction and then divided into ‘true’ (|Δt| < 42s) and ‘false’ (|Δt| > 42s) matches. The percentage of these matches is shown as a function of the window size in 1/*K_0_* that was applied to the matching. **b.** The gain in specificity by using ion mobility as a function of the window size in 1/*K_0_*.

### Feature coverage and label-free Quantification

The main reason for applying matching between runs is to make the MS1 features needed for quantification appear consistently across samples and by that remove the missing value problem that shotgun proteomics potentially possesses due to the stochasticity of MS/MS selection. We first assess the extent of the problem in technical replicates of cellular proteomes. For this we used data of ten replicates of HeLa cell lysate measured on the timsTOF pro as described in the Experimental Procedures and analyzed them with MaxQuant, once without and once with matching between runs. **Fig. 6a** shows for each of the ten technical replicates the number of protein groups identified and quantified without and with match between runs. While without matching between runs, on average 5358 protein groups were found per sample, match between runs increased this number by 381 on average. The number of protein groups that were quantified in all ten replicates is increased by matching between runs by nearly 1,000 to 5503 (**Fig. 6b**).

**Figure 6.**
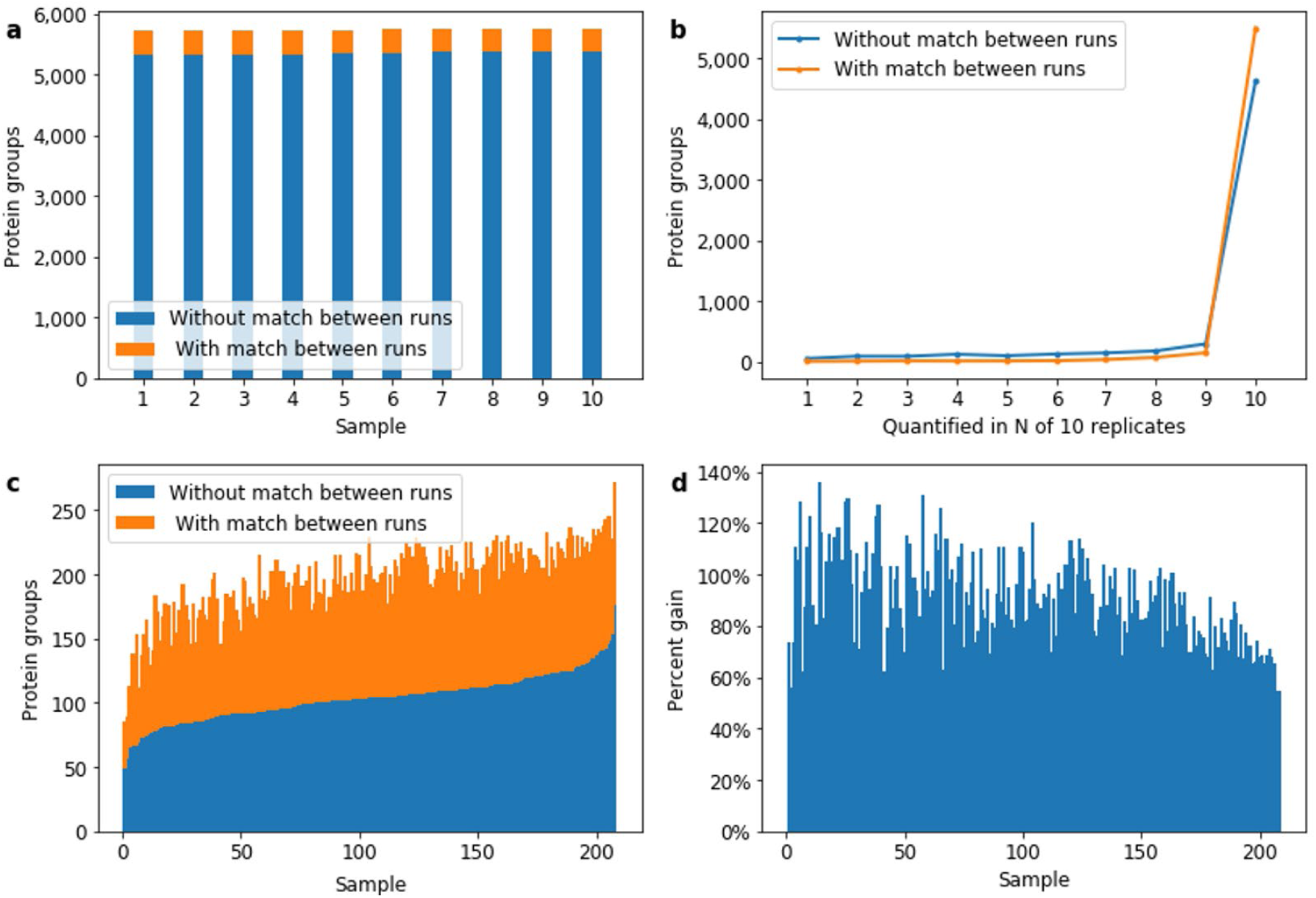
Protein quantification coverage. **a.** Number of protein groups quantified in 10 replicates without and with matching between runs. **b.** Number of proteins groups quantified in N out of 10 replicates without and with matching between runs. **c.** Number of protein groups quantified in 208 short human plasma runs. **d.** Gain of quantified protein groups in **c.** by matching between runs.

Reproducibility of quantification between samples is a much more challenging problem in plasma samples with their high dynamic range of concentrations and stronger variability. To this end we analyzed 208 plasma samples of human donors with single shot runs on a timsTOF pro with a 11.5 minutes LC gradient. (See experimental procedures.) In **Fig. 6c** the number of protein groups identified and quantified per sample is plotted for each of the 208 plasma samples in ascending order. Without matching between runs this number reaches from 49 to 176 protein groups with an average of 103. The gain in number of quantified protein groups achieved by matching between runs (**Fig. 6d**) is 90% on average.

To judge quantitative accuracy of label-free quantification, we recorded a dataset with known ground truth by mixing three cellular proteomes from different species in known ratios, similar to the strategy applied in the context of data-independent acquisition (DIA) data (29). While human proteins are expected to have a 1:1 ratio, all yeast proteins are expected to go up by a 2:1 ratio while all E. coli proteins go down by a 1:4 ratio. The MaxLFQ algorithm(20)can be applied to ion mobility enhanced data without major changes. Our adaptation to timsTOF data uses the signal intensities of the 4D MS1 features as input but is otherwise unaltered from its established LC-MS/MS version. The benchmark data consists of two replicate groups consisting of 3 samples each. The runs in a replicate group have been quantified as a single experiment with the MaxLFQ algorithm, which has been run once with and once without the matching between runs option activated. The resulting logarithmic LFQ intensities in the protein groups table have been normalized in the Perseus software by subtracting the most frequent value. Fold changes between the two replicate groups are plotted against the summed LFQ intensity in **Fig. 7a** and **b** without and with matching, respectively. The respective fold change data distributions are shown in **Fig. 7c** and **d**. The fold changes are centered on the positions that were expected for the three different species. The number of quantified proteins increases from 5819 to 6626 by using the matching between runs. There is no discernible intensity-dependent trend in the fold changes, meaning there are no nonlinearities in quantification at the upper or lower end of the dynamic range. Also the matching does not lead to a systematic change in the population of fold changes or to a discernible increased rate of wrongly quantified proteins, supporting the view that the matching is specific, as independently found with the method introduced in **Fig. 5**.

**Figure 7.**
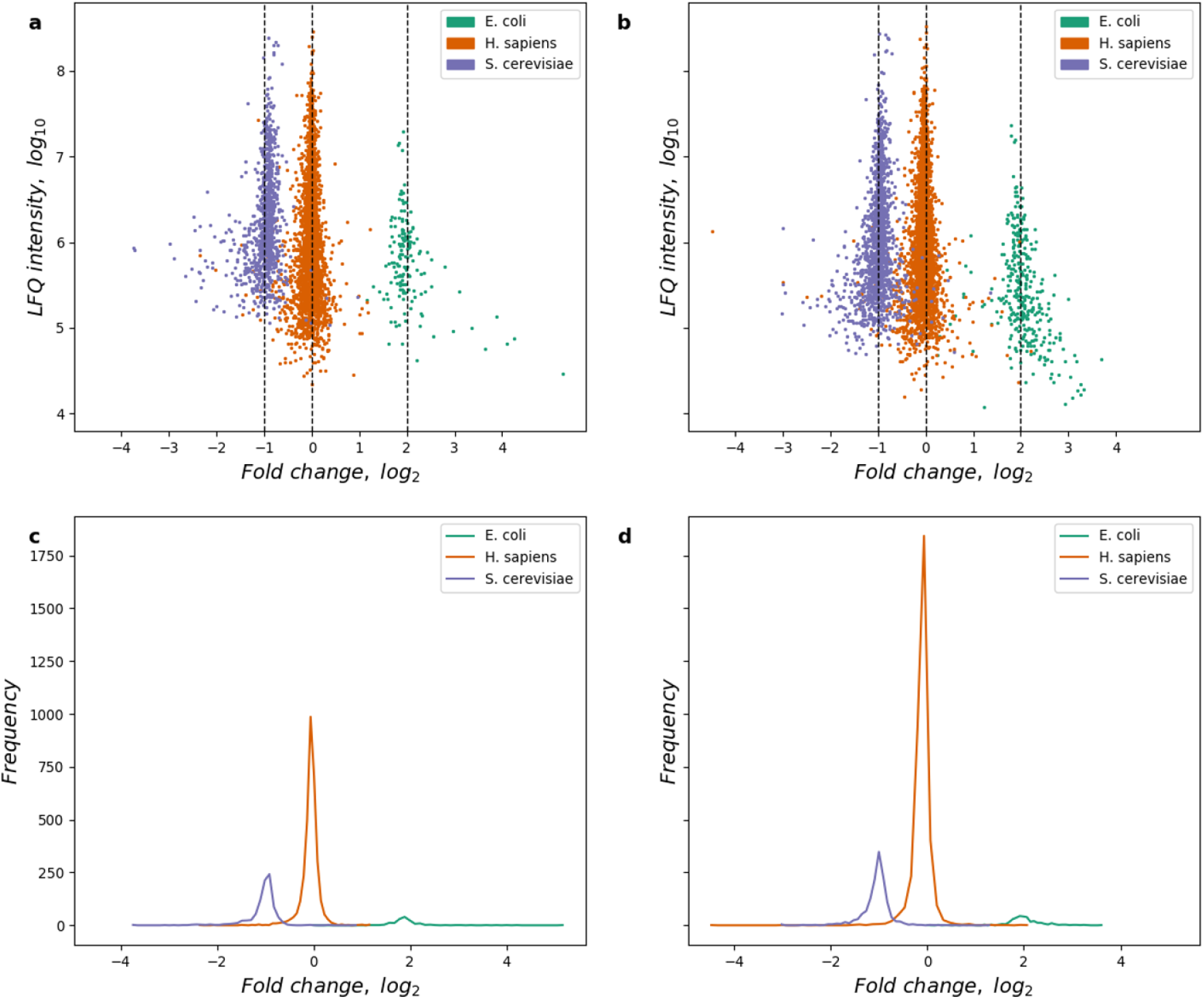
LFQ on a benchmark dataset. **a.** LFQ intensity plotted against fold change between replicate groups. (Both logarithmic.) Vertical lines correspond to fold changes expected by the mixing of species-derived samples. **b.** Same as **a**, but with matching between runs. **c.-d.** Histograms of data in **a-b** projected on the horizontal axes.

### Computational performance

A major challenge in the project was to achieve acceptable computational performance for large datasets with many LC-IMS-MS/MS runs being analyzed together in one project. Parallelization on multiple cores was achieved for nearly the entire computational workflow. Performance improvements will continue to be worked on and integrated into future software versions. The current split of computation time for a typical dataset onto the different workflow components can be seen in **Table 1**. Dominating parts are feature detection, MS/MS preparation, the searches, as well as the alignment.

## DISCUSSION

We implemented a novel computational workflow in MaxQuant that uses peptide CCS values to largely alleviate the missing value problem, typical in DDA shotgun proteomics. The newly developed MaxQuant CCS re-alignment algorithm does not require external or recognizable calibrants added to the sample, and successfully aligns ion mobilities from different runs against each other, prior to performing MBR. It has profound implications on protein quantification which we show to be possible with label free methods to good proteome depths and with enhanced precision. The combination of this particular IMS-QTOF hardware and the MaxQuant software provides the user with a robust platform for shotgun proteomics with deep and reproducible proteome coverage. Furthermore, the missing value problem of DDA shotgun proteomics due to its inherent stochasticity is appreciably reduced.

Currently, one of our development emphases is to improve computational performance of the total workflow. While with the current performance, data analysis is feasible and not prohibitively slower compared to conventional LC-MS/MS data, speed gains in future software releases will still have a positive impact on the everyday usage of MaxQuant on standard computer hardware. We are in an iterative development process of finding computational bottlenecks in the workflow and alleviating them. Given the significant improvements described in this paper, we foresee a CCS aware MaxQuant version soon that will run just as fast as the current version for conventional LC-MS/MS data. Going forward, the combination of DIA with IMS in combination with MaxQuant’s advanced feature detection and recalibration routines has the potential to increase robustness of proteomics analyses even further. The application of deep learning for the prediction of MS/MS spectra, retention time and ion mobilities, and its future integration into MaxQuant (30) has the potential to further improve protein identification rates and LFQ.

## ACKNOWLEDGEMENTS

We acknowledge Nicolai Bache and Ole Bjeld Horning from Evosep Biosystems for their support to run the plasma proteome samples and Thomas Kosinski from Bruker for sample preparation and LC-MS/MS analyses.

