## Supplementary material for "MaxQuant software for ion mobility enhanced shotgun proteomics"

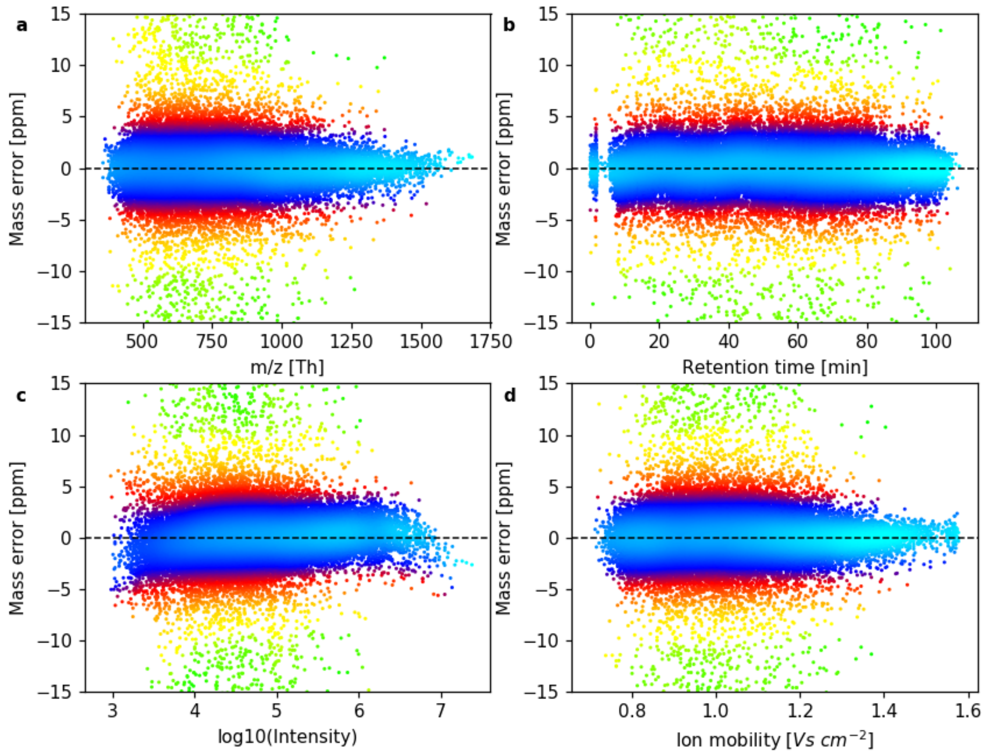


**Supplementary Figure 1. Mass error after recalibration**. **a.-d.** Residual mass errors after complete recalibration, showing the dependence of the residual mass error on *m*/*z* (**a.**), retention time (**b.**), logarithm of the peak intensity (**c.**) and ion mobility (**d.**). Colors reflect the density of data points.
